# Cycad coralloid roots contain bacterial communities including cyanobacteria and *Caulobacter* spp that encode niche-specific biosynthetic gene clusters

**DOI:** 10.1101/121160

**Authors:** Karina Gutiérrez-García, Edder D. Bustos-Díaz, José Antonio Corona-Gómez, Hilda E. Ramos-Aboites, Nelly Sélem-Mojica, Pablo Cruz-Morales, Miguel A. Pérez-Farrera, Francisco Barona-Gómez, Angélica Cibrián-Jaramillo

**Affiliations:** Evolution of Metabolic Diversity, Unidad de Genómica Avanzada (Langebio), Cinvestav-IPN, Km 9.6 Libramiento Norte, Carretera Irapuato-León, CP 36821, Irapuato, Guanajuato, México; Ecological and Evolutionary Genomics Laboratories, Unidad de Genómica Avanzada (Langebio), Cinvestav-IPN, Km 9.6 Libramiento Norte, Carretera Irapuato-León, CP 36821, Irapuato, Guanajuato, México; Escuela de Biología, Universidad de Ciencias y Artes del Estado de Chiapas, Libramiento Norte Poniente s/n, Col. Lajas-Maciel, CP 29029, Tuxtla Gutiérrez, Chiapas, México

**Keywords:** Cycads, coralloid roots, *Dioon*, cyanobacteria, *Caulobacter*, EcoMining

## Abstract

Cycads are the only early seed plants that have evolved a specialized root to host endophytic bacteria that fix nitrogen. To provide evolutionary and functional insights into this million-year old symbiosis, we investigate endophytic bacterial sub-communities isolated from coralloid roots of species from *Dioon* (Zamiaceae) sampled from their natural habitats. We employed a sub-community co-culture experimental strategy to reveal both predominant and rare bacteria, which were characterized using phylogenomics and detailed metabolic annotation. Diazotrophic plant endophytes, including *Bradyrhizobium, Burkholderia, Mesorhizobium, Nostoc*, and *Rhizobium* species, dominated the epiphyte-free sub-communities. Draft genomes of six cyanobacteria species were obtained after shotgun metagenomics of selected sub-communities and used for whole-genome inferences that suggest two *Dioon-*specific monophyletic groups and a level of specialization characteristic of co-evolved symbiotic relationships. In agreement with this, the genomes of these cyanobacteria were found to encode unique biosynthetic gene clusters, predicted to direct the synthesis of specialized metabolites, mainly involving peptides. After combining genome mining with metabolite profiling using multiphoton excitation fluorescence microscopy, we also show that *Caulobacter* species co-exist with cyanobacteria, and may interact with them by means of a novel indigoidine-like specialized metabolite. We provide an unprecedented view of the composition of the cycad coralloid root, including phylogenetic and functional patterns mediated by specialized metabolites that may be important for the evolution of ancient symbiotic adaptations.

## Introduction

Cycads (Cycadales) are gymnosperms of Permian origin that developed a specialized root organ to host symbiotic bacteria. Cycad lateral roots can develop into coralloid roots, a dichotomous and coral-like small cluster of roots, typically growing above ground, that acquire symbiotic bacteria (Norstog et al. 1997). Its main function seems to be nitrogen fixation for the plant (Bergersen et al. 1965, Groobelar et al 1986), similar to the adaptive functions in legume nodules, but having appeared millions of years before them. In natural habitats coralloid roots appear in early life stages of the plant (Halliday et al. 1976); and in adults mainly in habitats with poor or inaccessible nutrients, such as sand dunes, sclerophyll forests, steep rock outcrops with high exposure to salt, and lowland forests with recurrent fires (Grove et al. 1980; our observations). Cycads are currently among the most endangered plant groups mostly due to habitat loss and poaching, and given that it is possible that coralloid roots and their bacteria are a key early trait that enabled cycads to thrive and adapt to changing environments during millions of years, their study becomes relevant.

Coralloid root endophytes have been studied since the 19th century [(Grilli 1980) and references therein]. However, most studies have focused on resolving the taxonomy of cultured cyanobacteria, which can be visually detected in the root forming a green ‘cyanobacterial ring’ under differentiated cortical cells lying beneath the epidermis (Storey 1968). These studies have been carried out with samples collected from either botanic garden collections in greenhouses (Zimmerman et al. 1992, Costa et al. 1999, Thajuddin et al. 2010), natural populations (Yamada et al. 2012, Cuddy et al. 2012), or a mixture of both (Costa et al. 2004, Gehringer et al. 2010, Zheng et al. 2018). Studies testing for the specificity of cyanobacteria report contrasting results regarding the specialization of coralloid root symbionts. In wild populations a single *Nostoc* strain was found in different coralloid roots (Costa et al, 2004). Other studies report a significant degree of biodiversity (Grilli 1980, Ow et al. 1999, Yamada et al. 2012). There is also inconclusive information on the origin and transmission of cycad bacterial endophytes (Cuddy et al. 2012, Lobakoba et al 2003a).

Anatomical studies of the cyanobacteria ring have shown the presence of mucilaginous or protein-rich material, which may host other associated bacteria (Grilli 1980, Baulina et al. 2003, Lobakova et al. 2003b). The taxonomic associations of these bacteria, however, have been mostly suggested on the basis of generally low-resolution markers (Grobbelaar et al. 1987, Huang et al. 1989, Thajuddin et al. 2010, Zvyagintsev et al. 2010, Zheng et al. 2018), and doubts about their endophytic nature remain to be dissipated. In addition to nitrogen fixation there have been suggestions of other unknown roles for the coralloid root (Grilli 1980), but there is no clear evidence of a broader function to date. Likewise, various chemical, physical and physiological processes appear to regulate the cycad-bacteria interaction (Lobakova et al. 2003a, Meeks 2009), although genes coding for these specialized mechanisms of the symbiosis have been not identified. The presence of unique specialized metabolites in the cycad coralloid root bacterial endophytes is of interest to us because they may be a result of co-evolution between the cycad host and the endophyte bacterial community (Scherlach, et al. 2018, Warshan et al. 2018).

Certain soil-dwelling and aquatic bacteria, including cyanobacteria, have genomic plasticity (Warshan, et al., 2018) and the capacity to synthesize specialized metabolites with overwhelming chemical diversity, believed to be produced to cope with biotic and abiotic pressures. These bacterial traits are concomitant with large genomes that code for specialized metabolites in rapidly evolving genetic units called biosynthetic gene clusters (BGCs) of about 25-40 Kbp (Tan, 2007). Specialized metabolites play an important role in competition and collaboration in microbe-microbe and host-microbe interactions, mainly acting as molecular signals (Massalha et al 2017). Hence, the term of ‘specialized metabolite’ rather than secondary metabolite or natural product provides a more accurate description of the evolution and biology related to these compounds.

Our goal in this study is to investigate the endophyte bacterial community of the coralloid roots of *Dioon* species, including *D. merolae* De Luca, Sabato & Vázq.Torres, *D. caputoi* De Luca, Sabato & Vázq.Torres and *D. edule* Lindl., which are long-lived, entomophilous, dioecious, and arborescent cycads endemic to Mexico (De Luca et al. 1981, Lázaro-Zermeño et al. 2012). The subject species of this study, *D. caputoi, D. edule* and *D. merole*, are smaller species with shorter trunks, fronds, and cones (Norstog et al. 1997), and belong to two closely related phylogenetic clades (Gutiérrez-Ortega et al. 2017). We collected coralloid root samples from natural populations in different habitats from their natural range, currently distributed in moderate population sizes of a few hundreds of individuals throughout Queretaro, Puebla, Oaxaca and Chiapas, in Central and South of Mexico. We provide evidence of the existence of a diverse endophytic bacterial community inside the coralloid root beyond cyanobacteria, which consists of a core of diazotrophic bacteria, and the remarkably small Alphaproteobacteria *Caulobacter*. Unique BGCs in monophyletic Nostocales and *Caulobacter* endophytes suggest functional specificity throughout the evolutionary history of the cycad-cyanobacteria coralloid root system, and that some of these metabolites may mediate bacterial interactions within this symbiotic niche.

## Methods

### Overall strategy

To overcome technical difficulties in characterizing the breadth of microbial diversity in these environmental samples, we used EcoMining, a combined sub-community co-culture, metagenomics and phylogenomics strategy to detect and measure taxonomic diversity and phylogenetic relationships in the endophytes of the cycad coralloid root (Cibrián-Jaramillo and Barona-Gómez 2016) (Fig. 1). We grew and isolated bacteria from environmental samples using a diverse set of semi-selective media that aim to capture as much of the cultured bacterial diversity (*t0*). Simultaneously, we enriched the same samples for specific sub-communities in co-cultures grown under oligotrophic conditions specific for cyanobacteria, namely, BG-11^0^ medium. This second approach aims to capture other bacterial groups that interact with, and depend upon, the metabolic capabilities of cyanobacteria grown under oligotrophic conditions. The sub-community co-cultures were grown over time and sampled after one month (*t1*) and at the end of one year (*t2*), leading to bacterial isolates, their genomes and metagenomes. Specifically, six cyanobacterial draft genomes from nearly axenic cultures (strains JP106B, JP106C, TVP09, RF15115, 35k25 and 33k59), and two draft genomes reconstructed from metagenomic data (RF31YmG and 3335mG) were obtained. These genomic sequences were mined for BGCs potentially directing the synthesis of specialized metabolites. Also, two *Caulobacter* strains, D5 and D4A, associated with cyanobionts from *D. edule* were isolated, and their genomes sequenced and mined for niche-specific metabolites. Both cyanobacteria and *Caulobacter* strains were characterized after metabolite profiling using multiphoton excitation fluorescence microscopy. Specific methods, materials and analytical tools are described in the following subsections.

**Figure 1.**
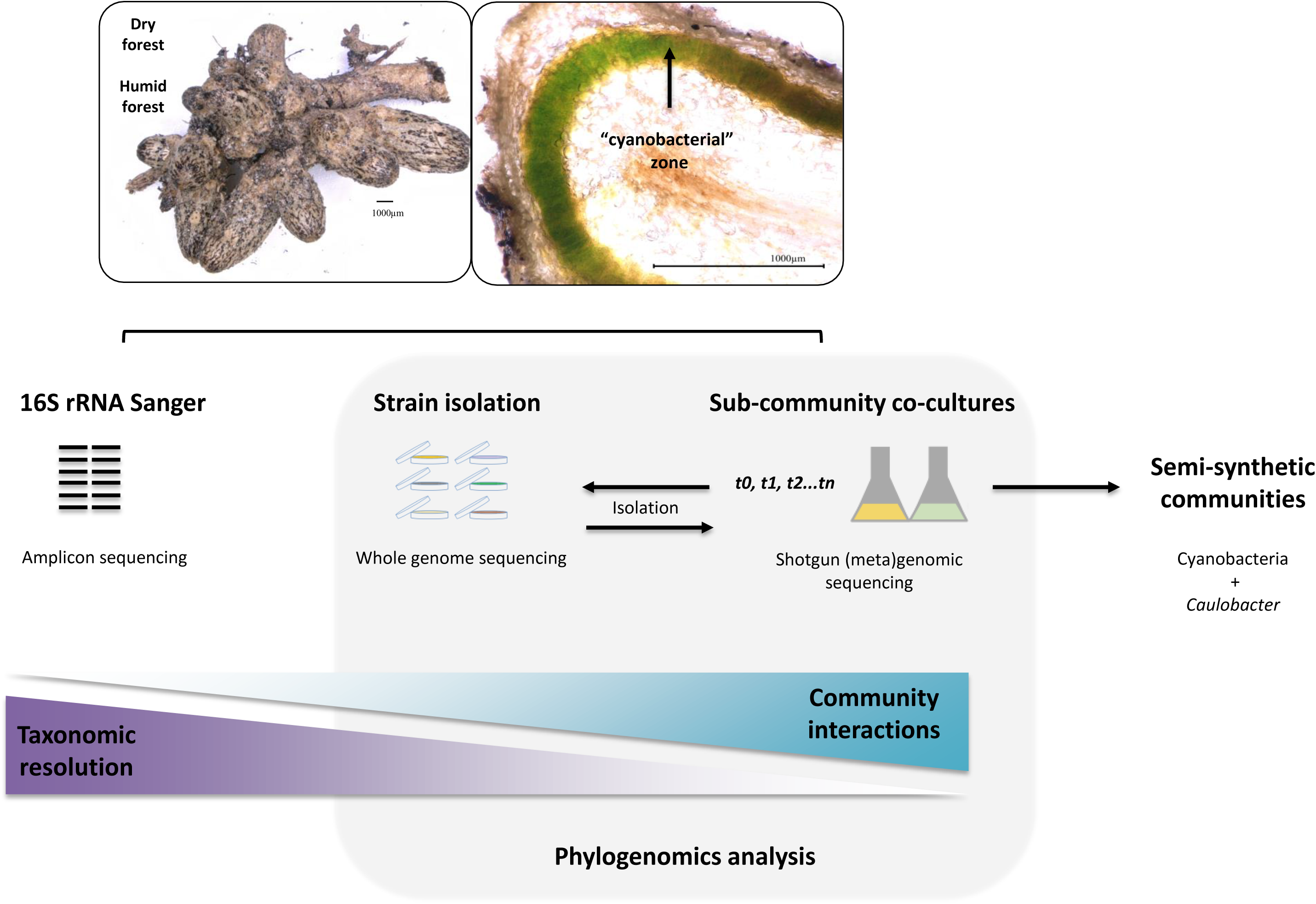
Diagram of experimental strategy to capture bacterial diversity (EcoMining). Coralloid roots from cycads growing naturally in deciduous tropical forests were sampled. Endophytes from the macerated root were isolated, following two strategies: directly from the sample (*t0*) and after enrichment of sub-communities co-cultures in BG-11^0^ (*t1, t2*). In both cases, cultivable bacteria were obtained using an array of six different media. Co-cultures were characterized using shotgun metagenomics, and the resulting data were used to select representative genomes from the endophyte culture collection that we mined for functional and evolutionary information.

### Field collections

We sampled coralloid roots from wild populations for each of the three *Dioon* species based on their previously reported distribution (Gutiérrez-Ortega et al. 2017). We sampled from two *D. merolae* populations in deciduous tropical forests, at JP Oaxaca Mexico (JP) at 560m above mean sea level, with an average annual precipitation of 320 mm and average annual temperature of 18°C; and at RF, Oaxaca, Mexico (RF) at 900m above mean sea level, with 2500 mm and 25 °C annual average precipitation and temperature, respectively. We also sampled one population for *D. caputoi* in Xeric shrubland, Tehuacan Valley Puebla, in Puebla, Mexico (TVP), at 1600m above mean sea level with an average annual precipitation of 474 mm and average annual temperature of 18°C. We sampled one *D. edule* population in SJD, Queretaro, Mexico (SJD) at 1300m above mean sea level, with an average annual precipitation of 966 mm and average annual temperature of 24 °C. Exact coordinates and names of locations can be share upon request to preserve the integrity of these endangered populations.

In some cycads coralloid roots were easily visible above ground, while in others we dug to about 20 to 30 cm around the main root until coralloid roots were found, especially when they were adult plants. We generally found 6 to 12 plants with coralloid roots, almost exclusively in seedlings, or in adult plants living on rock outcrops. A total of 20 coralloid apogeotropic roots from all species were removed from plants that appeared healthy, to minimize the negative impact on the plant. Roots were placed in 15 ml sterile Falcon tubes (Beckton Dickinson) and transported immediately to the laboratory at room temperature.

### Coralloid root processing

We focused our sequencing effort on 13 samples: i) two *D. merolae* samples from two plants from JP, named JP2 and JP6, and two *D. merolae* samples from two individuals from RF, named RF1 and RF3; ii) one sample from *D. caputoi* (TVP09); and iii) eight samples from four individuals from SJD, named SJD1 to SJD8. The remaining coralloid root samples were stored at −80 °C for subsequent studies. We treated the coralloid root in a laminar flow hood (Nuaire Model Nu-126-400) with a series of washes to remove exogenous bacteria from the rhizosphere or epiphytic contamination. Each root was introduced in 50 ml sterile Falcon tubes containing 10 ml of each of the following solutions, and gently stirred for: three minutes in hydrogen peroxide (H^2^O^2^), seven minutes in 70% ethanol, 30 seconds in sterile dd-MilliQ water, four minutes in 6% sodium hypochlorite (NaClO), and three one-minute washes in sterile dd-MilliQ water. After this procedure, we plated out water from the last wash in two media described below. Lack of growth in the last wash was considered a negative control. Only samples complying with this criterion were used for endophyte isolation (i.e. *Caulobacter*). We undertook two approaches to bacterial isolation (Fig. 1): sampling without enrichment directly from field samples after treatment for removal of epiphytes (*t0*), and sampling from the enriched co-cultures (*t1*).

### Bacterial growth and isolation in selective media

To isolate bacteria from the original epiphyte-free field samples (*t0*) and after enrichment (*t1*), coralloid roots were macerated and the macerate or co-culture broth were used as inoculant. In all cases, roots were macerated in 10 ml of sterile water using a pestle and mortar until plant material was completely disintegrated. After enrichment (*t1*) we recover bacteria expected to be initially present in low abundances and required time to grow, and that did so as a response to the sub-community nutritional interactions (e.g. amino acids derived from the process of fixing nitrogen). We used 100 µl from the root macerate to directly isolate bacteria in Petri dishes containing a total of six different media, chosen to selectively (oligotrophic, four media) or non-selectively (eutrophic, two media) recover bacterial diversity as much as possible. The four selective media used were chosen to target bacterial groups that are known to be either plant endophytes or rhizosphere bacteria, and included: 1) *Caulobacter* medium (glucose: 1 g/L; peptone: 1g/L; yeast extract: 1.5 g/L; trace metals solution: 1 mL/L; and 10 g/L of agar for solid medium); 2) *Rhizobium* medium (mannitol: 10 g/L; dipotassium phosphate: 0.5 g/L; magnesium sulfate: 0.2 g/L; yeast extract: 1 g/L; sodium chloride: 0.1 g/L; final pH 6.8; and 20 g/L for solid medium; 3) ISP4 and ISP4^N-^, for isolation of actinomycetes (starch: 10.0 g/L; dipotassium phosphate: 1 g/L; magnesium sulfate: 1 g/L; sodium chloride: 1 g/L; ammonium sulfate: 2 g/L or none in the case of ISP4^N-^; calcium carbonate: 2 g/L; ferrous sulfate: 1 mg/L; magnesium chloride: 1 mg/L; zinc sulfate: 1 mg/L; final pH 7.0; and 20 g/L for solid media); 4) BG-11^0^, a cyanobacteria medium (sodium nitrate: 1.5 g/L; dipotassium phosphate: 0.04 g/L; magnesium sulfate: 0.075 g/L; calcium chloride: 0.036 g/L; citric acid: 0.006 g/L; ferric ammonium citrate: 0.006 g/L; EDTA (disodium salt): 0.001 g/L; sodium carbonate: 0.02 g/L; final pH 7.1 and agar solid media 10.0 gr/L. The non-selective, rich media, included: 5) Nutrient Broth (BD Bioxon, Mexico); and 6) As in *Caulobacter* medium, but supplemented with mannitol (*Caulobacter* + mannitol medium): 1g/L, with aim of providing a carbon source closer to that hypothetically encountered inside the cycad root.

### Bacterial growth and cultivation in sub-communities

100 µl of the macerated roots free from epiphytes from the 13 samples (two *D. merolae* samples from JP, two *D. merolae* samples from RF, one from *D. caputoi*, and eight *D. edule* samples from SJD), which passed the negative growth control were used to inoculate 100 ml of media in 250 ml flasks. The remaining macerated roots not used for fresh cultures were kept as frozen stocks for future studies (−80 °C in 50% glycerol), although community viability after freezing was found to diminish over time. We used selective oligotrophic medium, i.e. BG-11^0^ (medium No. 4) for these experiments. BG-11^0^ cyanobacteria-centric co-cultures were grown for up to one year with constant stirring, with cycles of 16/8 hours of light/darkness. Sampling of the oligotrophic sub-communities was carried out at two different times, between 30 to 45 days (*t1*) and one year (*t2*): *D. merolae* oligotrophic sub-communities were sampled after 30 days (*t1*) and one year (*t2*), and treated independently; whereas *D. edule* co-cultures were sampled after 45 days (*t1*) and *D. caputoi* after 35 days (*t1*), although the latter did not include bacterial isolation. Bacterial isolates were only obtained from *t0* and *t1*, whereas shotgun metagenomics were obtained for *t1* (all three species) and *t2* (solely *D. merolae*).

### Genomics and shotgun metagenomics

The genomic sequence of five cyanobacteria isolates, JP106B, JP106C, RF15115, 33k59 and 35k25, were obtained from samples JP2, JP6 and RF1 (*D. merolae*) plus SJD1 and SJD2 (*D. edule*), respectively (*t1*). The genome sequence of the cyanobacterial isolate TPV09 was obtained from coralloid roots of *D. caputoi*. After many passes in both solid and liquid BG-11^0^ medium, all the microorganisms were grown on BG-11^0^ plates before DNA extraction, which was done using a CTAB-phenol chloroform standard protocol. Metagenomic DNA of all enriched sub-community co-cultures (200 ml), coming from JP, RF and SJD samples, were extracted using the same protocol, with the exception that biomass was obtained from liquid cultures by high-speed centrifugation during 15 minutes. Additionally, two *Caulobacter* strains, D5 and D4A, were isolated from 35k25 sample (*t1*) using ISP4 medium (medium No. 3). The axenic organisms were grown on liquid *Caulobacter* medium (medium No. 1) before DNA extraction, carried out using the protocol described previously.

DNA samples were processed with the truseq nano kit and sequenced at Langebio, Cinvestav (Irapuato, Mexico) using the MiSeq Illumina platform in the 2X250 paired end reads format (TVP09), and the NextSeq mid output 2X150 paired-end read format for all other samples. The reads for each library were filtered with fastQ and trimmed using Trimommatic version 0.32 (Bolger et al. 2014). Assembly of JP106B, JP106C, 33k59, 35k25, RF15115, TVP09, D4A and D5 sequences was done *de novo*, whereas the RF31YmG and 3335mG genomic sequences were obtained from metagenomic-assembled reads of the sub-community co-culture RF31Y of *D. merolae* (*t2*), and 33k59 plus 35k25 of *D. edule* (*t1*), respectively. For these assemblies, the newly obtained sequence of strain JP106C and *Nostoc punctiforme* PCC 73102 were used as references. The metagenomic reads were filtered by mapping them against the assembly of each reference genomes with BWA (Li et al. 2009). In all cases the resulting reads were assembled with Velvet (Zerbino et al. 2008) using different *k*-mers: the assemblies with the largest length and the smaller number of contigs were selected and annotated using RAST (Aziz et al. 2008). The genomic sequences that were released as independent metagenomic-assembled genomes (from 33k50 plus 35k25, RF31Y, TVP09 and JP6106C, respectively, labeled in Figure 2) are 3335mG (GenBank PRJNA472998), RF31YmG (GenBank PRJNA360315), T09 (GenBank PRJNA360300) and 106C (GenBank PRJNA360305).

**Figure 2.**
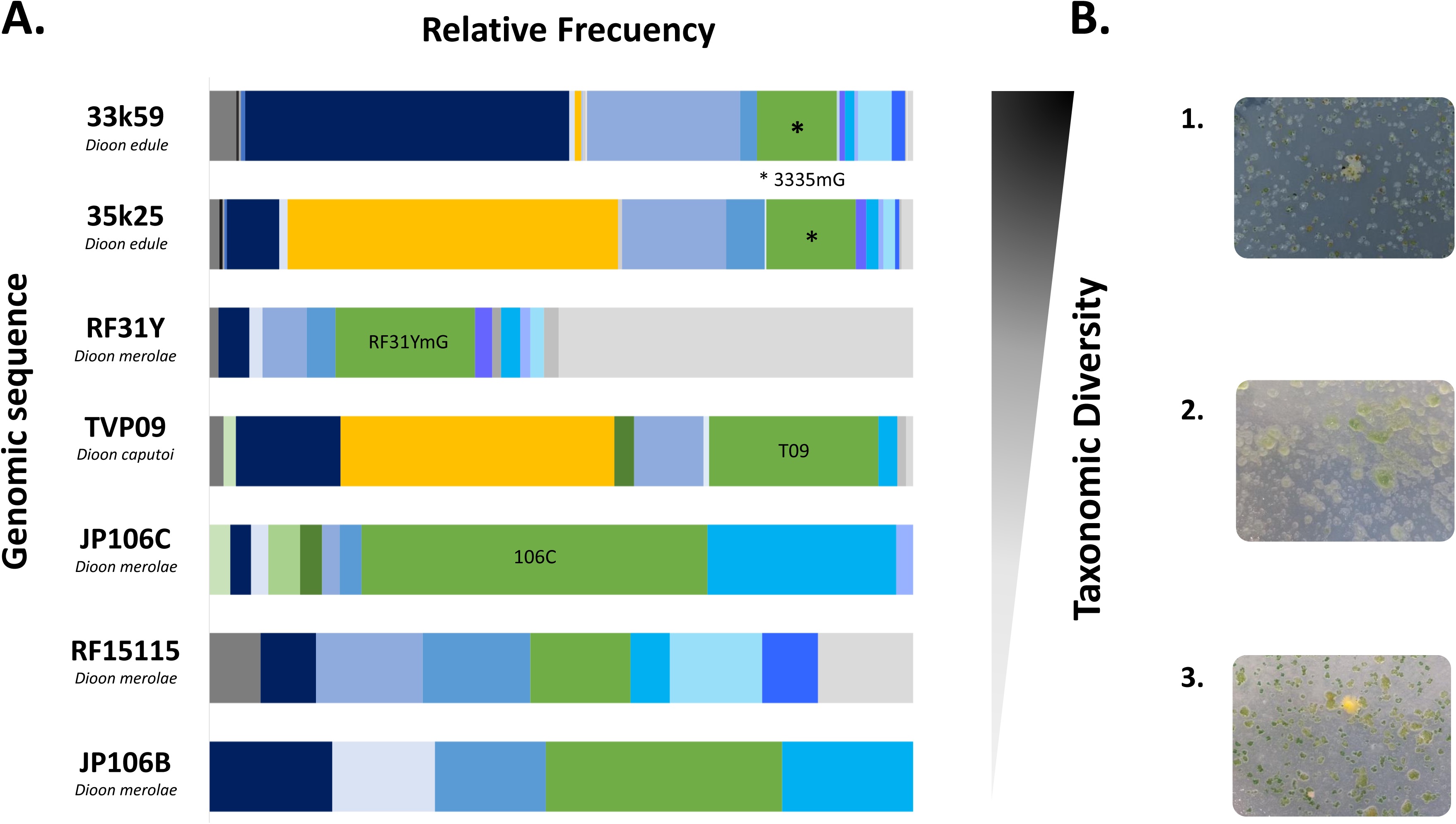
Taxonomic diversity of selected endophytic sub-community co-cultures characterized with shotgun metagenomics. **A.** Seven endophytic genomic sequences obtained from sub-community co-cultures were organized according to their taxonomic diversity, which includes an enrichment of diazotrophic organisms: cyanobacterial species (marked in green) and *Bradyrhizobium, Burkholderia, Mesorhizobium* and *Rhizobium* species (marked in blue). In some samples *Caulobacter* was also found (marked in yellow). 3335mG, RF31YmG, T09 and 106C assemblies were obtained from 33k50 plus 35k25, RF31Y, TVP09 and JP6106C, respectively. See Methods for more details of the sequences nomenclature. **B.** Selected examples of the endophytic sub-communities, grown in solid-medium as co-cultures: 1. high diversity, 2. medium diversity and 3. low diversity.

To confirm the presence of diazotrophic organisms in our sub-community co-cultures, we searched for the NifH protein sequence, an enzyme involved in nitrogen fixation and relevant in symbiotic relationships (Warshan et al 2018). We used the NifH protein from *Nostoc punctiforme* PCC 73102 (B2IV87), *Bradyrhizobium japonicum* OX=375 (I4DDG5) and *Mesorhizobium japonicum* LMG 29417 (Q98AP7) from the NCBI databases (July 2018).

### Taxonomic diversity

We estimated taxonomic diversity using the 16S rRNA gene sequences after PCR amplification and Sanger sequencing for our entire bacterial endophyte collection (*t0* and *t1*). PCR products of 1.4 Kbp in length, obtained using the F27 and R1492 primers (Lane 1991), were cloned and sequenced using the Sanger method. Taxonomic identification was made using Blastn with an initial cut-off e-value of 1e-5 against the SILVA database (Quast et al. 2013) and more than 98% sequence identity. We used the phylogenetic position of the top 10 hits from each search without duplicated matches, to determine both taxonomic diversity and phylogenetic relationships. We only report genus-level taxonomy as species determination can be biased by reference 16S rRNA databases.

We used Kraken (http://ccb.jhu.edu/software/kraken/) to assign taxonomic labels to metagenomic DNA sequences based on exact alignment of *k*-mers (Wood et al. 2014). Kraken was used to exclude sequence contaminants from the draft assembly, allowing us to generate a symbiotic cyanobacteria marker database as reference for future classification. We made a standard Kraken database using NCBI taxonomic information for all bacteria, as well as the bacterial, archaeal and viral complete genomes in RefSeq (October 2016). This database contains a mapping of every *k*-mer in Kraken's genomic library to the lowest common ancestor in a taxonomic tree of all genomes that contain that *k*-mer. We summarized the results in genera-level tables for each metagenome and filtered taxonomy hits that had 50 or more reads assigned directly to a taxon. To visualize this diversity we selected the best genomic sequences base on two criteria: i) Coverage > 25x, and ii) Assemblies with less of 20,000 contigs. From these, we chose those assigned to each taxa with ≥ 500 reads.

### Reconstruction of phylogenomic relationships

To reconstruct a complete phylogeny of the cyanobacteria and *Caulobacter* strains using their whole genomes, we adopted a phylogenomics approach that we termed OrthoCores based on our in-house scripts (https://github.com/nselem/orthocore/wiki, dockerhub: nselem/orthocores). We assume that single gene or even housekeeping multilocus phylogenies are often not enough to resolve the evolutionary history of certain taxonomic groups, and thus total evidence tree evidence approaches must be used (Narechania et al. 2012). We first obtained a set of orthologous groups that are shared among all high-quality genomes, or the ‘core genome’, excluding paralogs and signatures of expansions (not all genes from a gene family are part of the Bidirectional Best Hits, or BBH set). We define orthologs from the core genome as hits that satisfy that all gene members are BBHs, all versus all, to cyanobacteria and *Caulobacter*. To place our samples within the cyanobacteria and *Caulobacter* phylogeny, we used OrthoCores to generate the tree. For the cyanobacterial phylogenetic tree, we selected closed and well-annotated genomes, and for Nostocales all the publicly available genomes. We included representatives from public Chroococcidiopsidales, Oscillatoriales, Synechococcales, Chroococcales, Pleurocapsales, Gloeobacterales and Nostocales and the eight genomic sequences obtained in this study. The *Caulobacter* phylogenetic tree was constructed with 22 publicly available genomes including our two *Caulobacter* genomes (D5: GenBank PRJNA472998 and D4A: GenBank PRJNA472998) and *Hyphomonas jannaschiana* VP2, used as outgroup.

The OrthoCores cut-off e-value was 1e-6, although most genes had a 1e-50 score given our all *versus* all criterion. Set of sequences were concatenated and aligned using MUSCLE v3.8.31 with default parameters (Edgar 2004), and trimmed by Gblocks (Talavera et al. 2007) with five positions as minimum block length, 10 as a maximum number of contiguous non-conserved positions, and only considering positions with a gap in less than 50% of the sequences in the final alignment. The final cyanobacterial matrix included 81 taxa and 198 core protein sequences with 45475 amino acids that represent a set of 71 cyanobacterial genomes including our six cyanobacterial genomic sequences (table s1), plus the RF31YmG and 3335mG metagenomic assemblies, and *Gloeobacter kilaueensis* JS1 and *Gloeobacter violaceus* PCC 7421 as outgroups. To build each cyanobacteria subsystems tree, the core proteome was used and proteins were classified using RAST (Aziz et al. 2008) and were used to construct a concatenated matrix. Each concatenated matrix was used to construct a phylogenetic tree using the methodology described above. The final *Caulobacter* matrix included 82 core protein sequences with 20902 amino acids (table s2). The resulting matrices were used to reconstruct the trees with MrBayes, using a mixed substitution model based on posterior probabilities (aamodel[Wag]1.000) for proteins for one million generations. Both trees were visualized with FigTree http://tree.bio.ed.ac.uk/software/figtree/).

### Genome mining for BGCs

To identify BGCs potentially directing the synthesis of specialized metabolites among selected cyanobacteria and *Caulobacter*, we annotated our genomic sequences, as well as selected genomes from the database, using antiSMASH (Weber et al. 2015). The predicted BGCs in both our cyanobacteria and *Caulobacter* strains (248 and 7 potential BGCs, respectively) were manually curated (only 49 cyanobacterial antiSMASH outputs did not pass this filter), leading to 77 and 7 non-redundant BGCs used as references for subsequent searches in all of the Nostocales and *Caulobacter* selected genomes, respectively. This process allowed us to identify both conserved and specific BGCs throughout our databases generated from publicly available genomes on NCBI (table s3 and s4). We considered that a BGC hit was found whenever any given enzyme, plus any other enzyme gene from the cognate BGC, were found in a vicinity of 20 genes, using a blast e-value cut-off of 0.001.

The prediction of the chemical structures of the putative specialized metabolites associated with some BGCs, namely, those *Dioon-*specific present in our six cyanobacterial genomic sequences (BGCs 1, 5 and 12), and a BGC specific within our *Caulobacter* strains, were done after domain identification and specificity prediction of adenylation and acyl transfer domains, using the non-ribosomal peptide synthetase (NRPS) and polyketide synthases (PKS) NRPS-PKS server (Bachmann and Ravel 2009), PRISM (Skinnider et al. 2015) and antiSMASH (Weber et al. 2015). The functional inferences of selected BGCs present in these strains, in parallel with their structural chemical prediction, was carried out after in-depth functional annotation of all genes belonging to these BGCs, using standard databases and sequence similarity approaches, together with a detailed revision of the literature.

### Metabolite profiling using multiphotonic microscopy

To probe if the genus *Caulobacter* co-exists with our cyanobacterial strains, we exploited the occurrence of specialized metabolites produced by these microorganisms. We obtained metabolite profiles of both isolated microbial cultures using intensity, time, and wavelength-resolved multiphoton excitation fluorescence microscopy, with a LSM880-NLO Multiphoton Microscope (Zeiss, Germany) coupled to an Infrared laser Ti: Sapphire (Chameleon vision II, COHERENT, Scotland). To perform *in vitro* assays, we used one of our *Caulobacter* endophytes, D5 strain, and our cyanobacteria isolate 35k25. Each isolate was grown on different plates containing BG-11^0^ medium. We also cultured both isolates together on plates containing BG-11^0^ and BG-11 media. All the samples were visualized through multiphoton excitation fluorescence microscopy, adding sterile water as a propagation medium. The observation areas were made using the objective 60x/1.3 NA ∞-0.17 (Zeiss Plan-Neofluar, Germany). The samples were excited at different excitation energy (700-850nm). Three emissions were detected: i) Multialkali detector (Ch1), using to collect photons between 371 to 471nm; ii) Detector GAsP (Ch2), using to collect photons between 501 to 601nm, and iii) Detector Multialkali (Ch3), using to collect photons between 616 to 688nm. Scans were performed in the image acquisition mode and in the lambda mode to obtain the spectral fingerprint of each analyzed area. The spectral signature is specific for every molecule and determined for each pixel of scanned image. This data was used for the subsequent digital separation of fluorescent molecules.

## Results

### *Dioon* coralloid roots show diazotrophic endophyte diversity beyond and within cyanobacteria

The 16S rRNA Sanger sequences that we obtained yielded 267 isolates distributed in 19 families and 12 bacterial orders, with 29 genera in total, representing most of the known bacterial groups. 86% of the taxa identified can be taxonomically classified as diazotrophic plant endophytes, including species belonging to the genera *Bradyrhizobium, Burkholderia, Mesorhizobium, Nostoc*, and *Rhizobium* (table s5). We also isolated *Bacillus*, which was previously reported as associated with the outside of the coralloid root (Lobakova et al. 2003a); *Streptomyces*, previously isolated as an epiphyte (Zvyagintsev 2010); and four strains identified as *Nostoc* spp, this genus previously identified in several cycads. However, as it will be further discussed in detail in the subsection dealing with the genomic analysis, these isolates turned out to be a micro-community resistant to axenic cultivation. Moreover, we continuously isolated *Caulobacter*, a genus that has been associated with cyanobacteria from *Azolla* ferns (Newton and Herman, 1979) and aquatic cyanobacteria (Bunt, 1961, Berg et al. 2008), but to our knowledge this is the first report of *Caulobacter* as a cycad endophyte.

Metagenomic sequencing of endophytic sub-communities was successful for JP2, JP6, RF1, RF3 and SJD1 to SJD8 grown until *t1*, and RF31Y and JP61Y, grown until *t2*. Basic indicators of metagenome quality are provided in table s6. For taxonomic analysis of these metagenomic data we focused on the most abundant genera (≥ 50 reads assignment of each taxa). These results are reported as the total number of taxa present in relation to which is the most abundant genus among these taxa, reported as a percentage, as follows: RF1, 24/*Xanthomonas* (40%); RF3, 21/*Xanthomonas* (63%); JP2, 27/*Nostoc* (24%); JP6 24/Xanthomonas (28%); SDJ1, 32/*Mesorhizobium* (41%); SDJ2, 36/*Bradyrhizobium* (24%); SDJ3, 36/*Caulobacter* (23%); SDJ4, 35/*Mesorhizobium* (31%); SDJ5, 37/ *Mesorhizobium* (41%); SDJ6, 50/*Bradyrhizobium* (25%); SDJ7, 24/*Bradyrhizobium* (34%) and SDJ8, 31/*Caulobacter* (35%). As expected, we observed different bacterial proportions for the *t2* samples, RF31Y, 42/*Nostoc* (55%) and JP61Y, 44/*Xhantomonas* (36%), when compared with their cognate counterparts from *t1*. The presence of *Nostoc* in both *t1* and *t2* samples was also confirmed (table s7).

To represent the taxonomic diversity of our endophytic sub-communities, shown as solid-medium co-cultures in Figure 2, we selected only the genomic sequences that passed our criteria: 33k59, 35k25, RF31Y, TVP09, JP106C, RF15115 and JP106B (see also Fig. S1).

Our endophytic sub-communities are enriched with diazotrophic organisms, including cyanobacteria belonging to the Nostocales order, such as those from the genera *Nostoc* (Thajuddin et al. 2010, Yamada et al. 2012) and *Calothrix*, previously reported in *Encephalartos* (Grobbelaar et al. 1987, Huang et al. 1989, Gehringer et al. 2010) and in *Cycas revoluta* (Thajuddin et al. 2010). Other diazotrophic organisms, such as *Bradyrhizobium, Burkholderia, Mesorhizobium*, and *Rhizobium* species, were also found. Moreover, *Caulobacter* species were found in 33k59, 35k25 and TVP09 samples (Fig. 2). The presence of diazotrophic organisms in our sub-community cultures was confirmed by Blast-positive hits of the NifH enzyme to *Bradyrhizobium* and *Mesorhizobium* species with 99% sequence identity.

Overall, the 16S rRNA Sanger results and the taxonomic analysis of our metagenomic data agrees with the bacterial diversity reported using single-amplicon metagenomics of environmental samples (Suarez-Moo et al. 2018). Among the most abundant, we confirm the occurrence of *Amycolaptopsis, Burkholderia, Caulobacter, Microbacterium, Pseudomonas, Ralstonia, Rhizobium, Serratia, Sphingobium, Sphingomonas, Streptomyces* and *Xanthomonas*, in addition to the Nostocales genera, as *Dioon* coralloid root endophytes.

### Cyanobacteria endophytes are a monophyletic group within the Nostocales

Given that cyanobacteria phylogenetic history is likely reticulated (Zhaxybayeva et al. 2006), we adopted a whole-genome inference approach to explore the phylogenetic position of our genomic sequences, including JP106B, JP106C, RF15115, 33k59, 35k25, and TVP09, and two draft genomes reconstructed from metagenomic data of co-culture sub-communities, RF31YmG and 3335mG. We reconstructed a phylogeny from 198 proteins identified as the core proteome encoded in 75 selected genomes including our own sequences, which provide a broad taxonomic coverage and represent good quality sequences (table s1, table s3). With the exception of the Nostocales, we found lack of monophyly of the cyanobacteria groups in our phylogeny (Fig. S2). However, our genomic sequences TVP09 from *D. caputoi*, plus JP106B, JP106C and RF31YmG from *D. merolae*, form a monophyletic clade, and group with *Tolypothrix* sp. PCC 7601. As can be seen in Figure 3, their clade is sister to *Calothrix* sp. PCC 7507 and *Fortiea contorta* PCC 7126, both reported as aquatic organisms. These organisms have not been tested for symbiotic competence, although a related species, *Calothrix rhizosoleniae*, is a symbiont of the diatom *Chaetoceros* (Foster et al. 2010). Likewise, RF15115 from *D. merolae*, with 33k59, 35k25 and 3335mG from *D. edule*, form a monophyletic clade with *Nostoc punctiforme* PCC 73102 from the cycad *Macrozamia* and *Nostoc* sp. KVJ20 from the liverwort *Blasia pusilla* (Warshan et al. 2018).

**Figure 3.**
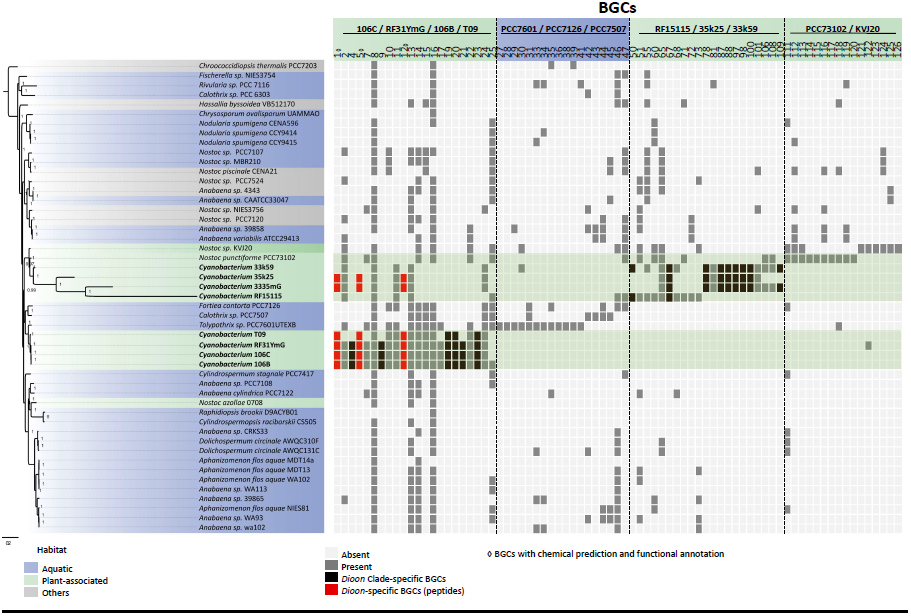
Phylogenomic tree of selected Nostocales species and their Niche-specific BGCs. Tree reconstructed with 198 conserved proteins to place our eight cyanobacterial genomic sequences, 106B, 106C, T09, RF31YmG plus RF15115, 33k59, 35k25 and 3335mG. Habitats for each species are indicated with colored bullets. The closely related organisms showing a sign of metabolic specialization, are highlighted with a green box. BGCs that are *Dioon* clade-specific (black boxes) and *Dioon*-specific (red boxes), with chemical structure prediction and functional annotation (◊) are shown.

In agreement with our phylogenetic reconstruction, our genomic sequences were notably syntenic with the genome of KVJ20, indicating that these organisms may share adaptive traits within equivalent ecological niches, i.e. specialized organs devoted to nitrogen fixation. It should be noted that aside from *Nostoc punctiforme* PCC 73102, the genome of a cyanobacterial strain isolated from the coralloid roots of a *Cycas revoluta* plant, termed *Nostoc cycadae* WK-1 (Kanesaki et al. 2018), was recently released. WK-1 strain, isolated and kept under laboratory conditions for almost four decades before its sequencing, was not included in our analyses as it does not group with the symbiotic cyanobacteria reported herein (Gagunashvili and Andresson, 2018). This latter report deals with cyanolichens genomes, but includes some of our genomic sequences previously released.

Long branches of 3335mG and RF15115, suggesting divergence from their closely related organisms, are also obvious. To further characterize this observation, trees based on gene sets from the core proteome, which correspond to 16 metabolic subsystems, were reconstructed, as different phylogenetic signatures between them may contribute to long branches. All of the subsystems trees showed divergent branches for both 3335mG and RF15115, albeit with similar topologies, with only two exceptions: nitrogen metabolism, consisting or only one regulatory protein, and virulence disease and defense (Fig. s3). Thus, even in the absence of a pending detailed taxonomic description of our cyanobacterial isolates, we can conclude that (i) these cycad endophytes belong to the nitrogen-fixing endophytic Nostocales clade; (ii) they are closely related despite their different *Dioon* host species; and (iii) within *Dioon*, there are two phylogenetic groups that include organisms that may be evolving relatively fast (Fig. 3). To gain functional insights from these evolutionary patterns we did an in-depth metabolic annotation as described next.

### Identification of BGCs in the sub-community metagenomes suggests metabolic specialization of *Dioon* cyanobacteria

Genome mining of specialized metabolites with an emphasis in our genomic sequences, plus five taxonomically closely related microorganisms identified in the previous section, unveiled a total of 248 antiSMASH hits (table s8). After manual curation of these outputs, a total of 77 non-redundant and well-defined BGCs (table s9) were used to analyze the distribution of these loci (Fig. 3). This analysis shows that most of these BGCs are uniquely present within plant-associated cyanobacterial genomes, including 43 BGCs contributed solely by the *Dioon-* specific cyanobacteria. Along these lines, we found that only four of these 77 BGCs are fully conserved BGCs present in all cyanobacteria. These include BGC 13, the well-characterized heterocyst glycolipid (Soriente et al. 1992); a ladderane lipids BGC (Sinninghe Damsté et al. 2002), termed BGC 16. And BGC 8 and 14, putatively directing the synthesis of a terpene and a bacteriocin-lantipeptide, respectively. The remaining BGCs could be overall annotated as five systems coding for NRPSs, of which some may contain additional PKSs (32%), bacteriocins (32%), terpenes (8%), lantipeptides (4%), and indoles (3%). Moreover, sixteen BGCs (21%) were classified as Others.

It should be noted that from the BGCs analyzed, 23% are only present in our cyanobacteria isolates, including three *Dioon*-specific BGCs (1, 5 and 12) and 15 *Dioon*-clade-specific BGCs (4, 9, 19, 20, 21, 23, 50,67, 78, 87, 88, 97, 98, 100 and 109), which are only present in cycads with coralloid roots and in the liverworth *Blasia* (Liaimer et al. 2016). Remarkably, organisms taxonomically closer to our strains, such as *Nostoc punctiforme* PCC7 3102 isolated from an Australian cycad of the genus *Macrozamia*, whose genome has been recently re-sequenced (Moraes et al. 2017), shows a reduced biosynthetic potential. Thus, we focus on the biosynthetic systems that are *Dioon*-specific, which have NRPS or PKS components, facilitating the prediction of their chemical structure. As mentioned, these include three BGCs exclusively found in our strains, i.e. BGCs 1, 5, and 12 (Fig. 4), which may provide examples of specialized metabolites with functional implications within the coralloid root, as further detailed.

**Figure 4.**
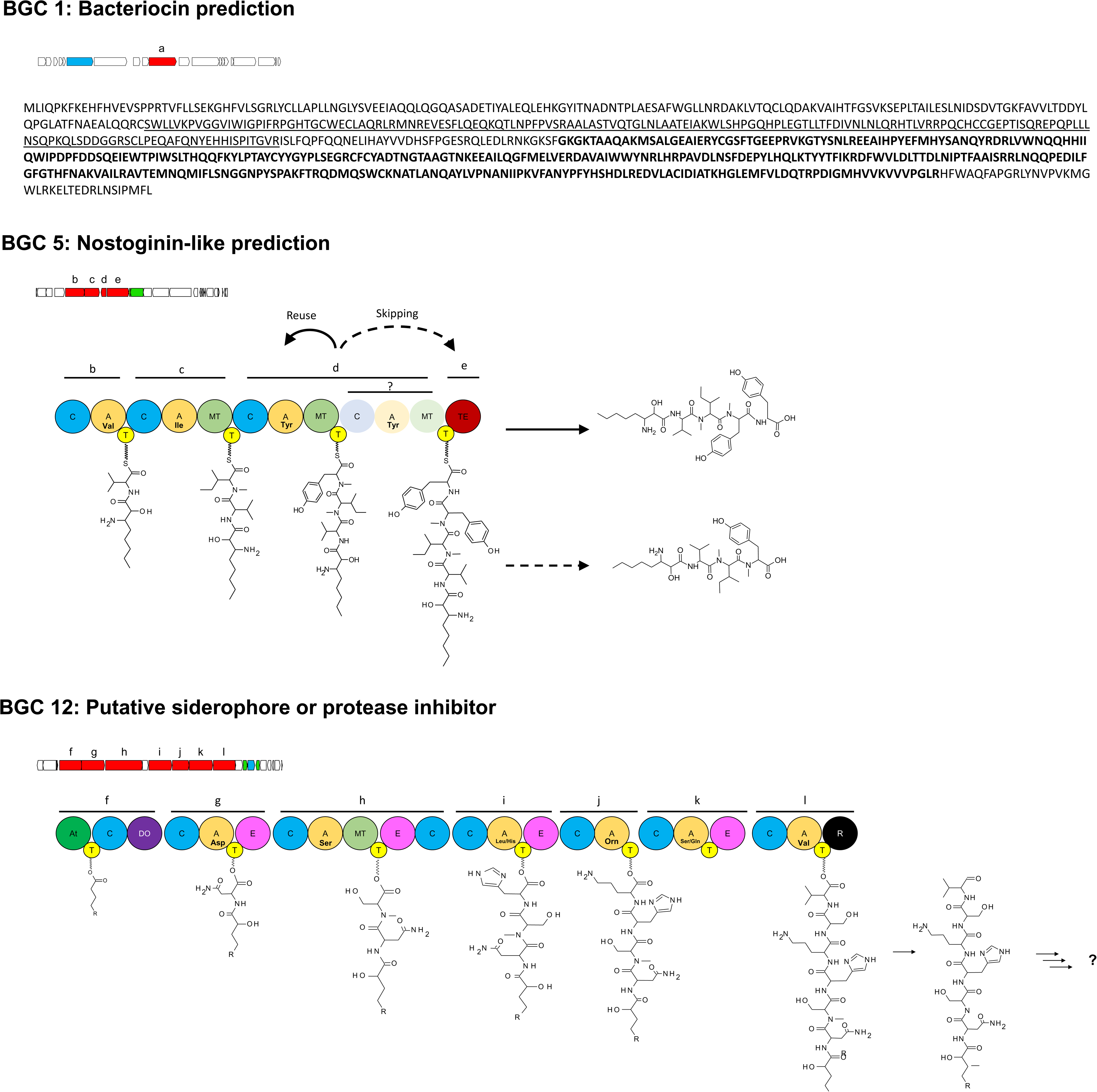
*Dioon*-specific BGCs with structure chemical prediction and functional annotation. Detailed analysis of BGCs 1, 5 and 12 present in the *Dioon-*specific strains is shown. Genes are shown as colored boxes: Core biosynthetic genes (Red), transport-related genes (blue), regulatory genes (green), and other genes (white). The tips of the boxes indicate the direction of their translation. Domain organization, biosynthetic logic and predicted products are indicated below each BGC: Acyltransferase domain (At), dioxygenase domain (DO), methyltransferase domain (MT), reductase domain (R). Underlined sequence: cyclodehydratase domain, black sequence: YcaO-like domain.

First, BGC 1, predicted to encode a bacteriocin, shares 53% of sequence identity with the biosynthetic genes of a bacteriocin isolated from the cyanobacteria *Pleurocapsa minor*. BGC 1 includes two key domains: i) the YcaO-like domain, involved in the peptidic biosynthesis of azoline, and ii) the cyclodehydratase domain, which could be a cyclodehydratase involved in the synthesis of a thiazole/oxazole-modified microcins (Dunbar et al., 2012). The sequence similarities detected are relevant as *Pleurocapsa minor* forms an intertwined community with *Calothrix* spp (Whitton 2012). Second, BGC 12 may code for an assembly line capable of synthesizing an N-terminal acylated hexapeptide with several modifications, such as the epimerization of four of its residues, the N-acylation of its second amidic bond, and the reduction of its C-terminal end to yield an aldehyde group. The N and C terminal modifications on this peptide would resemble small peptide aldehyde protease inhibitors, which have been previously reported on cyanobacteria (Fewer et al. 2013). Alternatively, the product of BGC 12 may be a siderophore, as iron-related genes were found next to the NRPS coding-genes, including a reductase domain implicated in the synthesis of iron chelators, e.g. myxochelin biosynthesis (Li et al. 2008).

Third, BGC 5, which codes for a NRPS system putatively directing the synthesis of a tripeptide consisting of valine, isoleucine and tyrosine residues. This prediction also includes informative modifications, such as N-terminal acylation, N-methylation at an amide bond of the isoleucine residue, and modification of the tyrosine residue by a domain of unknown function. A search for peptides containing the abovementioned modifications, performed with the server PRISM with the feature for de-replication of known chemical structures (Skinnider et al. 2015), directed our attention to nostoginins, specialized metabolites originally isolated from *Nostoc* spp (Ploutno et al. 2002). Interestingly, the biosynthetic pathway of this natural product remains to be identified, but it is known that nostoginins A and B, as well as their congeners microginins FR1 / SD755 and oscillaginins A / B (Sano and Kaya 1998), have protease inhibitory activity. In agreement with this observation, BGC 5 includes a peptidase-coding gene (Fig. 4).

It is clear from these results that the *Dioon* cyanobionts have distinctive metabolic capabilities, perhaps as a result of the co-evolution of these organisms within a diverse microbiome present in the coralloid root. In analogy with the pairwise interdependencies reported for *Pleurocapsa* and *Calothrix* (Whitton 2012), it is tempting to speculate about similar cases occurring within the *Dioon* coralloid roots microbiome. The predicted functions may be relevant for such microbial interactions, in addition to the interactions with its *Dioon* host species. As a first step towards the biochemical and functional characterization of our strains, with regards to the occurrence and role(s) of specialized metabolites, we focused in the cyanobacteria-*Caulobacter* putative interaction.

### Cyanobacteria-*Caulobacter* association may be mediated by specialized metabolites

During the course of our investigations, we consistently found *Caulobacter* species associated with cyanobacteria throughout our various methods, even when efforts to obtain completely axenic cyanobacterial cultures were carried out. In contrast, *Caulobacter* strains, whose genomes were sequenced from monocultures, could be isolated relatively easy. Two *Caulobacter* strains, D5 and D4A, were isolated from the sub-community co-cultures from *D. edule* (*t1*), which is the same that yielded the 35k35 genomic sequence. The genome sequences of these organisms shows similarity with *Caulobacter* sequences extracted from the genomic sequence 35k35. The *Caulobacter* phylogenomic tree, reconstructed using 82 proteins from 22 *Caulobacter* genomes and *Hyphomonas jannaschiana* VP2 as outgroup, places both of our strains together with *Caulobacter vibrioides* LNIY01. Little is known about this latter microorganism other than the fact that it was isolated from a selenium mining area, and that it may be capable of producing the phytohormone 3-indoleacetic acid (Wang et al. 2016).

After genome mining for the prediction of BGCs, as expected from an Alphaproteobacteria, the genus *Caulobacter* shows reduced biosynthetic potential, with only 90 BGCs from all 22 genomes. All the BGCs could be overall annotated as six systems coding for NRPSs, of which some may contain additional PKSs (4%), bacteriocins (37%), terpenes (20%), lassopeptides (19%), H-serlactone (14%), and fatty acid-saccharides (6%). Despite this limited metabolic diversity, however, we detected a unique BGC, classified as Other, that is present in our monophyletic clade but absent from all other *Caulobacter* genomes (Fig. 5A). This BGC, termed BGC 1, shows an overall 80% sequence similarity with respect to the indigoidine BGC, encoding a blue pigment described previously in various actinobacterial species, including *Streptomyces chromofuscus* (Yu et al., 2013), *Streptomyces aureofaciens* (Novakova et al. 2010), *Streptomyces lavendulae* (Kurniawan et al., 2014); but also in the distantly related Gammaproteobacteria, and plant pathogen, *Erwinia chrysanthemi* (Reverchon et al., 2002) and in the Alphaproteobacteria, and marine bacterium, *Phaeobacter* sp. strain Y4I (Cude et al., 2012).

**Figure 5.**
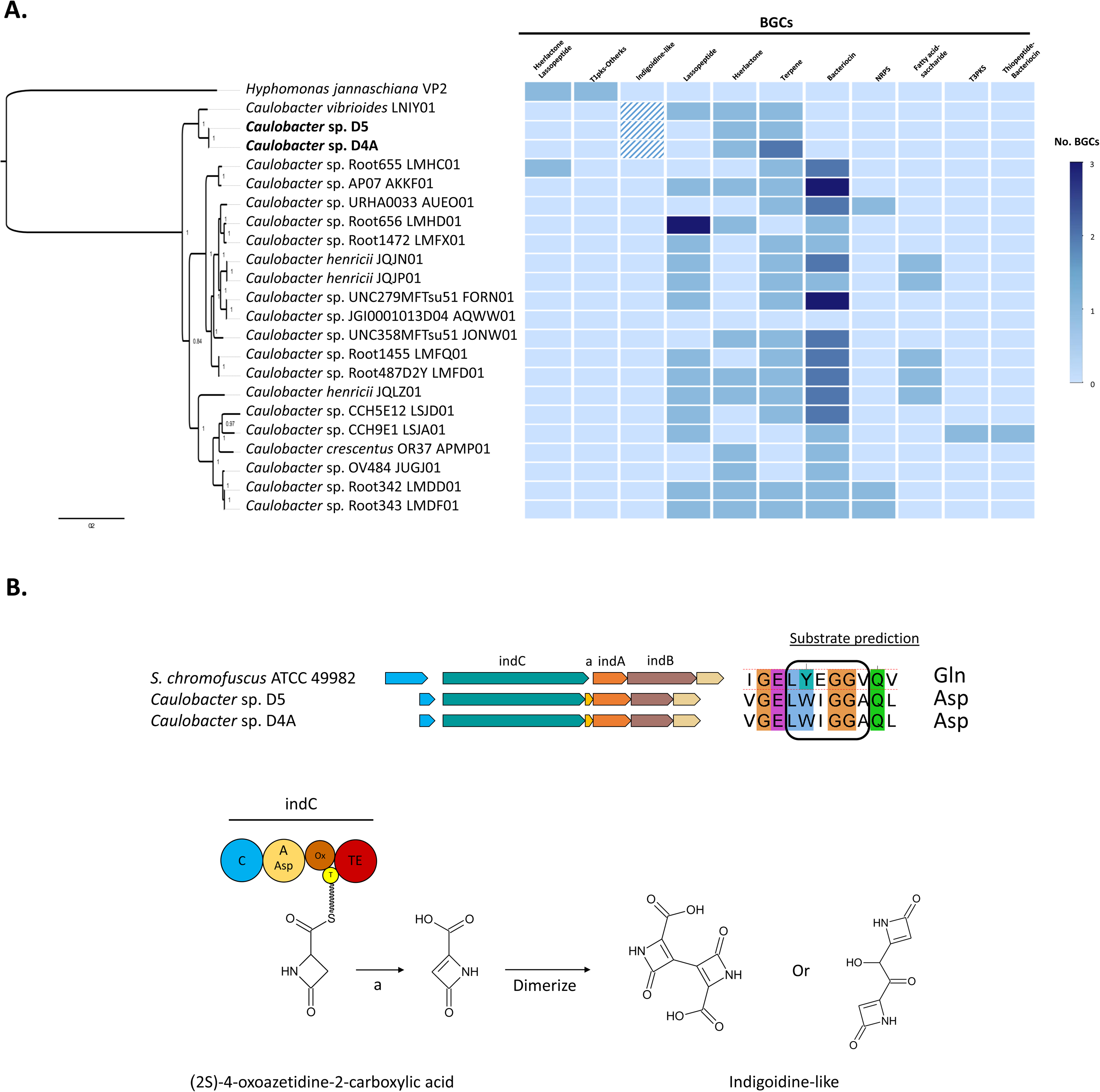
Phylogenomic tree of *Caulobacter* species and their BGCs distribution. **A.** Tree reconstructed with 82 conserved proteins to place our two *Caulobacter* strains, D5 and D4A, showing a sign of metabolic specialization. **B.** Chemical structure prediction and functional annotation of the aspartic acid-derived indigoidine-like metabolite.

Comparative analysis between BGC 1 and the indigoidine BGC shows similar organization, including four biosynthetic genes, one of which is an NRPS, two are transporter genes and one is a regulatory gene. A key difference was found in the adenylation domain of the NRPS, which instead of a glutamine residue as in *bona fide* indigoidine, we predicted it to recognize an aspartic acid (Reverchon et al., 2002) to form (2S)-4-oxoazetidine-2-carboxylic acid. Additionally, BGC 1 has a gene coding for an enoyl reductase domain, which may form a double bond in the oxoazetidine ring. This compound could dimerize to form an indigoidine-like compound (Fig. 5B). There are no reports of this predicted indigoidine-like metabolite in other bacteria, although synthesis of related congeners with antibacterial activity have been reported (O’Dowd et al., 2008). The distribution of the indigoidine BGC in divergent bacterial lineages, suggests horizontal gene transfer to fulfill biological functions yet-to-be determined.

To test if our *Caulobacter* endophytes co-exist with our cyanobacterial strains through indigoidine-like blue compounds, metabolite profiles based in multiphotonic microscopy were obtained (Fig. 6). For these experiments, microbial cultures of *Caulobacter* sp. D5 and Cyanobacteria sp. 35k25, both isolated from *D. edule*, were used. Three different emissions were detected, Ch1 (371-471nm), Ch2 (501-601nm) and Ch3 (616-688nm), after exciting samples at different excitation energies (700-850nm). To dissect these emissions we first observed the characteristic emission signals when each sample or strain was cultivated separately. The axenic culture of *Caulobacter* sp. D5 showed a unique and relevant metabolic emission between 550-570nm (green light). In contrast, for the Cyanobacteria sp. 33k25 sample, in agreement with the fact that it is a micro-community rather than a monoculture, three emissions were detected corresponding to chlorophyll (650-720nm, red light), the above mentioned *Caulobacter* species (550-570nm, green light), and a small signal located in the range of blue light (430-480nm, blue light). We suggest that later emission is related to the presence of the indigoidine-like compound predicted after genome mining.

**Figure 6.**
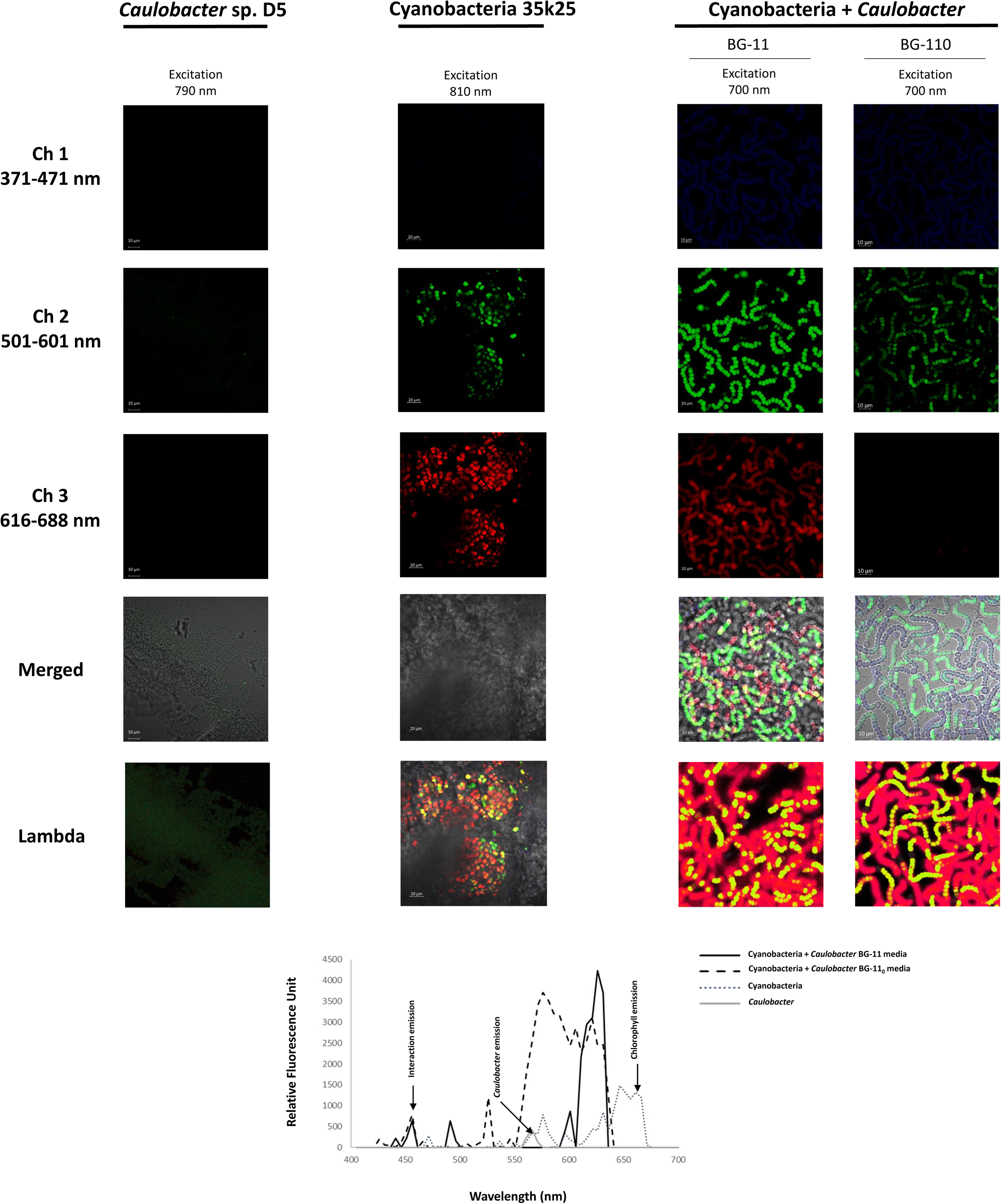
Metabolite profile *in vitro* of the Cyanobacteria-*Caulobacter* association. *Caulobacter* sp. D5 and Cyanobacteria sp. 33k25, in reality a micro-community, were grown independently on plates containing BG-11^0^ medium, and together on plates containing BG-11^0^ and BG-11 media. *Caulobacter* sp. D5, by its own, showed a unique and relevant metabolic emission between 550-570nm (green light). Cyanobacteria 33k25, showed three emissions corresponding to chlorophyll (650-720nm, red light), the *Caulobacter* species (550-570nm, green light) and a little signal proposed to be related to the predicted indigoidine-like metabolite (430-480nm, blue light). In both BG11 and BG-11^0^ synthetic bacterial co-cultures, three emission signals were detected corresponding to chlorophyll (650-720nm), the *Caulobacter* species (550-570nm), and remarkably, an increase of the signal related with the indigoidine-like metabolite (430-480nm). Refer to the text for further details.

To further investigate into the possible association between strains D5 and the cyanobacteria contained in 35k25, potentially mediated by the predicted indigoidine-like metabolite, *Caulobacter* sp. D5 and Cyanobacteria sp. 33k25 were cultivated together in plates containing BG11 and BG-11_0_ media. As expected from previous results, irrespective of the media, the synthetic bacterial culture emitted signals in the same wavelengths as chlorophyll (650-720nm), the *Caulobacter* species (550-570nm) and that suggested to be related to the indigoidine-like metabolite (430-480nm). Interestingly, the intensity recorded for each signal, in relative fluorescence units, was different for the two different media used. Specifically, the chlorophyll emission decreased in BG-11_0_, which may be due to the large number of heterocyst formed under nitrogen limitatuion. However, the emission intensity of the blue signal related to the indigoidine-like metabolite remained the same in both conditions, but became notably greater in the synthetic co-culture. This result suggests that the interaction between the cyanobacteria contained in 33k25 and the *Caulobacter* spp induces the production of the indigoidine-like metabolite.

## Discussion

Our evidence undoubtedly shows that there is a diverse bacterial community within the cycad coralloid root that includes all of the previously reported Nostocales, other taxa for which their endophytic origin and presence was unclear, and newly reported genera. This is congruent with recent reports (Zheng et al., 2018; Suarez-Moo et al., 2018), and with morphological studies observing mucilaginous material inside the coralloid root (Ow et al. 1999, Baulina et al. 2003). This supports previous studies in which different species of cycads host multiple shared Nostocales cyanobacteria (Yamada et al. 2012, Zimmerman et al. 1992, Zheng W et al. 2002, Gehringer et al. 2010), yet the monophyletic placement of our cyanobacterial samples clearly indicates a different evolutionary trajectory of *Nostoc* species associated with *Dioon* from their free-living counterparts. However, this phylogenetic separation is only clear when the whole genome is taken into account, as fewer genetic markers do not seem to have enough resolution to resolve the clade (Yamada et al. 2012). For instance, free-living *Nostoc* PCC 7120 grouped distantly to strains of symbiotic or plant-associated origin, while the also distant genome reported as *Nostoc azollae* is now taxonomically *Trichormus azollae* (Baker et al. 2003). Similarly, other non-cycad plant-associated *Nostoc* are more related to *Anabaena* and *Aphanizomenon*, congruent with previous reports (Papaefthimiou et al. 2008).

Our cyanobacterial endophytes are part of a clade that includes symbiotic species able to colonize specialized nitrogen-fixing cavities through plant tissues or inside coralloid roots. This observation is congruent with previous phylogenies that include hormogonia-producing symbiotic species as clustered together (Gagunashvili and Andrésson 2018, Warshan et al. 2018). The ability of different strains isolated from one plant species of infecting phylogenetically distant hosts, as previously reported (Johansson et al. 1994, Papaefthimiou et al. 2008, Whitton 2012), remains to be tested for our *Dioon* endophytes. We hypothesize that species within our phylogenetic clade could cross-infect different plant hosts, although their metabolic capacities and contributions to each cycad genus is very likely to change given their unique BGCs. This idea is consistent with previous cross-infection reports with the cycad symbiont *Nostoc punctiforme* PCC 73102 and the bryophyte *Anthoceros* (Enderlin and Meeks 1983), but goes deeper into possible mechanisms mediated by specialized metabolites.

Differences in genera identified with 16S rRNA and shotgun metagenomics of sub-communities are explained in part due to the biases in 16S rRNA markers, which can erroneously diagnose the taxonomic composition of endophytes in the cycad host (Louca et al., 2018). These differences can also be explained by other scenarios: i) rare groups present in low abundance can only be recovered in sub-community co-cultures on which they increase in biomass; ii) some organisms are fast growers irrespective of media, and will dominate in OTUs, simply by chance; iii) some groups are more media-specific; and/or iv) groups in BG-11^0^ (*t1*) are recovered as a result of functional interactions in cyanobacteria-associated groups. Also, technically speaking, a biased on the metagenomic data may be related to the DNA extraction protocol, which has been shown to confound diversity assessment of endophytic communities (Maropola et al. 2015).

The long-term one-year co-culture (*t2*) allowed us to explore at least some of these possibilities, as the initial amount of nitrogen available in these co-cultures became a limiting factor over time. Hence, the establishment of stable sub-communities after one year (with emerging and surviving taxa) suggests that autotrophic, nitrogen fixation and photosynthesis, may be a driving force in the assembly of the coralloid root community. With this observation in hand it is know possible to set out comparative and systematic analyses of the diazotrophic potential of these communities. As coupling the metagenomic analysis with traditional microbiology increased our ability to accurately identify interacting endophytes in the sub-communities, our adopted approach may help us to characterize microbial life strategies and their phylogeny in a complex system, which is an interesting challenge that has received increasing attention (Ho et al. 2017).

Notably, genome mining of the cyanobacterial strains revealed three BGCs unique to the *Dioon*-specific cyanobionts, in agreement with the hypothesis of a specialized community driven by metabolic adaptation regardless of the taxonomic plasticity. Of the three BGCs found in our isolates, BGC 12 and BGC 5 putatively coding for small peptide protease inhibitors, or a siderophore, could have a functional bearing on the evolution and biology of the *Dioon*-bacteria symbiosis. Peptide production may imply that proteolysis is involved in the cyanobacteria-cycad interaction, linked to the reconfiguration of the root architecture or the filtering of the microbiome, as it has been observed in arbuscular mycorrhiza and legumes (Takeda et al. 2007). A starting point to explore this possibility would be to confirm the predicted gene-metabolite relationship proposed here, by sequencing the genomes of nostoginin-producing organisms (Ploutno et al. 2002) together with their functional characterization. If a siderophore, BGC 12 could be mediating bacterial community interactions as previously reported (Cruz-Morales et al., 2017), and/or interactions with the host, adding to the growing notion that symbiotic relations occur under heavy influence of metabolic exchange.

The association between our *Caulobacter*, a small Alphaproteobacterium adapted to nutrient-poor conditions (Landt et al., 2010), and the *Dioon* cyanobacterial endophyte strains, is also remarkable. Both our two *Caulobacter* isolates are able to produce a specific and unique specialized metabolite, indigoidine-like compound, which could act as antimicrobial or phytohormone, potentially mediating the association between either organisms or their host. Hence, the *Caulobacter-*Nostocales consortium could be playing a previously unknown role inside the coralloid roots of cycads by producing few but specific natural products, making them specialized metabolites. The metabolite profiling of this system using multiphonic microscopy, a novel approach in many ways, opens the possibility of looking into these presumed interactions *in planta*, and warrants more detailed chemical analysis of the metabolic exchange occurring within the microbiome of the cycad coralloid root.

In sum, our study demonstrates the broad taxonomic bacterial diversity of the coralloid root, which combined with monophyletic Nostocales with at least three unique BGCs and an associated *Caulobacter* with a unique BGC, could be a sign of the constraints imposed by the transition from a free-living lifestyle, perhaps initially even as a bacterial consortium outside the plant, into a symbiotic lifestyle inside a plant’s specialized organ. As data is gathered from more genomes of bacterial cycad symbionts isolated from different environments of the rhizosphere-coralloid root ecological niche, it will be possible to test for deeper co-evolutionary relationships. Genomes from other endophytic genera identified here, could inform the evolutionary and ecological nature of cycad-bacterial and bacterial-bacterial interactions, which in turn could direct detailed functional characterization of these ancient plant specialized communities.

## Acknowledgements

This work was supported by CONACyT #169701 and FON.INST./265/2016 (SA-FC2015-2-901) to ACJ. CONACyT #179290 and #177568 to FBG. We acknowledge Juan Palacios and Antonio Hernández for help during field collections, as well as Flor Zamudio for technical support.

## Data deposition

The draft genomes generated during the current study are available in the GenBank public repository as follows:

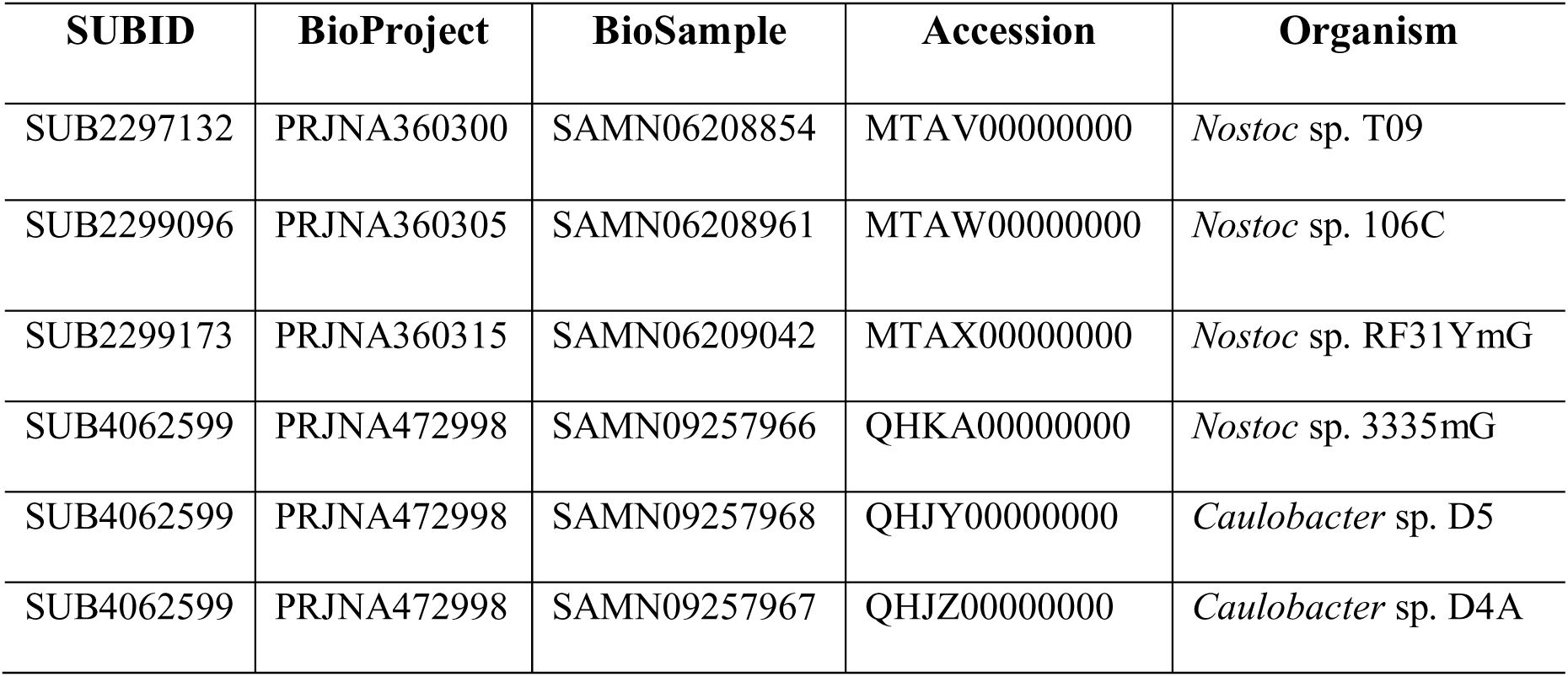

Metagenomes are available at:

***Dioon merolae* coralloid roots.** https://www.ncbi.nlm.nih.gov/bioproject/369422
***Dioon edule* coralloid roots.** http://www.ncbi.nlm.nih.gov/bioproject/472628
**Cyanobacterial genomic sequences.** https://www.ncbi.nlm.nih.gov/sra/SRP149992

## Supplementary files

Supplementary Tables1-9.xls

Table S1. Core proteome of cyanobacteria (198 proteins)

Table S2. Core proteome of *Caulobacter* (82 proteins)

Table S3. Cyanobacteria genomes used in this study

Table S4. *Caulobacter* genomes used in this study

Table S5. List of 271 isolated bacteria characterized with Sanger 16S rRNA

Table S6. Statistics and details of co-culture metagenomes sequenced

Table S7. Distribution and abundance of mOTUs from the co-culture metagenomes

Table S8. antiSMASH hits of each cyanobacterial genome.

Table S9. BGCs on the genomes of isolates JP106C, TVP09, JP106B, PCC7601, PCC7126, PCC7507, RF15115, 35k25, 33k59, PCC73102 and KVJ20

Supplementary Figures.pdf

Figure s1. Taxonomic diversity of selected endophytic sub-community co-cultures characterized with shotgun metagenomics

Figure s2. Phylogenomic tree of selected cyanobacterial genomes

Figure s3. Phylogenomic tree of selected cyanobacterial genomes base on sub-systems

